# Bidirectional restoration of sleep homeostasis in neurodegeneration via closed-loop auditory stimulation

**DOI:** 10.64898/2026.04.13.718146

**Authors:** Inês Dias, Marc A. Züst, Christian R. Baumann, Daniela Noain

**Author notes:** Corresponding author –; Address: Wagistrasse 14, 8952 Schlieren, Switzerland. Email addresses: Inês Dias; Marc Züst; Christian R. Baumann, Daniela Noain.

## Abstract

Sleep-wake disturbances in Alzheimer’s (AD) and Parkinson’s disease (PD) are linked to disrupted slow-wave activity (SWA) and SW-spindle coupling, critical for memory consolidation and protein clearance. While SWA enhancement shows promise in aging, disease-specific effects on SW synchronization (local, linked to sleep depth vs. global, linked to arousal), spindle dynamics and sleep homeostasis remain unexplored in neurodegeneration. Using mouse closed-loop auditory stimulation (mCLAS), we probed its acute effects on SW subtypes, spindle characteristics, SW-spindle coupling and sleep homeostasis in Tg2576 (AD) and M83 (PD) mice. mCLAS adaptively modulates SW subtypes and SW-spindle coupling depending on baseline impairments. In AD mice, it reduces global synchrony while potentiating local phenomena and network recruitment, restoring SW-spindle coupling strength and normalizing spindle power. In PD mice, mCLAS increases global synchrony, reorganizing 24-hour SW dynamics towards wild-type patterns, while reducing excessive SW-spindle coupling and enhancing spindle power. Critically, mCLAS bidirectionally regulates sleep homeostasis: boosting deficient slow-wave energy in AD and reducing excess in PD to restore physiological sleep. These findings suggest mCLAS acts via complementary mechanisms to rescue sleep homeostasis in neurodegeneration, by modulating cortical synchrony in AD and arousal in PD, offering a noninvasive path to potentially mitigate cognitive decline and pathological progression.

## Introduction

Sleep-wake disturbances are a common feature of both the asymptomatic preclinical and the clinical stages of Alzheimer’s (AD) and Parkinson’s disease (PD), with sleep believed to play a crucial role in the onset and progression of neurodegenerative processes^1–6^. Decreased slow-wave activity (SWA) and sleep continuity during non-rapid eye movement (NREM) sleep are believed to impact brain functions such as memory consolidation, synaptic renormalization and pathological protein clearance in patients and animal models of neurodegeneration^1,7–10^. Consequently, a two-way relationship between sleep and neurodegeneration is hypothesized, with altered sleep-wake patterns and neurodegeneration progression potentially worsened by NREM sleep disruption and vice versa^11–14^.

NREM sleep slow-waves (SW, <4 Hz) and spindles (10-15 Hz) are particularly affected by aging and neurodegeneration. Hallmarks include reduced SW and spindle density and amplitude, shallower slopes, longer SW durations, altered SW-spindle coupling with reverse spindle localization along the SW, and altered spindle duration and frequency^9,10,15–18^. Additionally, two distinctive SW synchronization processes co-exist during NREM sleep: global synchronization, lacking homeostatic decline, and local synchronization, which is homeostatically regulated^19–21^. These processes were reported to be altered in older adults^21^ but have not been thoroughly assessed in disease settings. Moreover, decreased SWA (0.5-4 Hz) could reflect altered local synchronization and impaired sleep dissipation, as SWA and slow-wave energy (SWE) are well-established correlates of sleep pressure dissipation and homeostatic regulation^22,23^.

Several SWA-induction protocols have enhanced NREM sleep in neurodegeneration, including pharmacological approaches in animal models^1,24,25^ and patients^26–28^, and non-pharmacological techniques like transcranial direct current stimulation and phase-targeted closed-loop auditory stimulation (CLAS) in older adults and mild cognitive impairment (MCI) patients, with improvements in memory recall and amyloid dynamics^29–36^. Despite this, the fundamental mechanisms underlying phase-targeted CLAS in neurodegeneration remain poorly understood, constraining its clinical translation ^37–39^; animal studies offer a unique opportunity to bridge this gap by enabling multilevel assessments. In this context, we recently established mouse CLAS (mCLAS) in two neurodegeneration models, demonstrating successful implementation in murine brain disease^40^.

However, whether SWA enhancement restores sleep homeostasis and not merely sleep quality in the context of neurodegeneration remains unexplored. Addressing this question is critical to determine whether CLAS-potentiated sleep recovery is mediated by homeostatic compensatory mechanisms in neurodegenerative diseases^40–42^. Here, we assessed mCLAS’ short-term effects on electrophysiological correlates of homeostatic sleep regulation in 7-month old Tg2576 transgenic mice, the AD line^43^ and 7-month old M83 transgenic mice, the PD line^44^. We characterized distinct SW synchronization processes, spindle dynamics, SW-spindle coupling and 24-hour SWE in AD and PD mice in comparison to age-matched healthy littermates (wildtype, WT). We found that mCLAS adaptively modulates these parameters in a disease-specific manner: in AD mice, it restored deficient sleep homeostasis by boosting SWE and normalizing SW-spindle coupling via enhanced local synchrony, whereas in PD mice, it reduced excessive SWE and excessive coupling while increasing global synchrony to reorganize 24-hour dynamics towards WT patterns. These results demonstrate that mCLAS bidirectionally rescues sleep homeostasis through complementary mechanisms targeting cortical synchrony in AD and arousal promoting phenomena in PD.

## Results

### SW dynamics are selectively modulated by mCLAS

To evaluate the modulatory effects of short-term mCLAS on SW synchronization processes, we implanted mice from the Tg2576 (AD) and M83 (PD) mouse lines, and their healthy littermates (WT controls), with EEG/EMG headpieces (**Figure 1a, b**). We classified SWs into types I (global synchrony), II (local synchrony), and III (intermediate processes) ^19–21^ (**Figure 1c**, see Methods), and quantified SW metrics including number, amplitude, slope and wavelength across experimental groups, protocol days and light/dark periods (**Figure 1d**).

**Figure 1.**
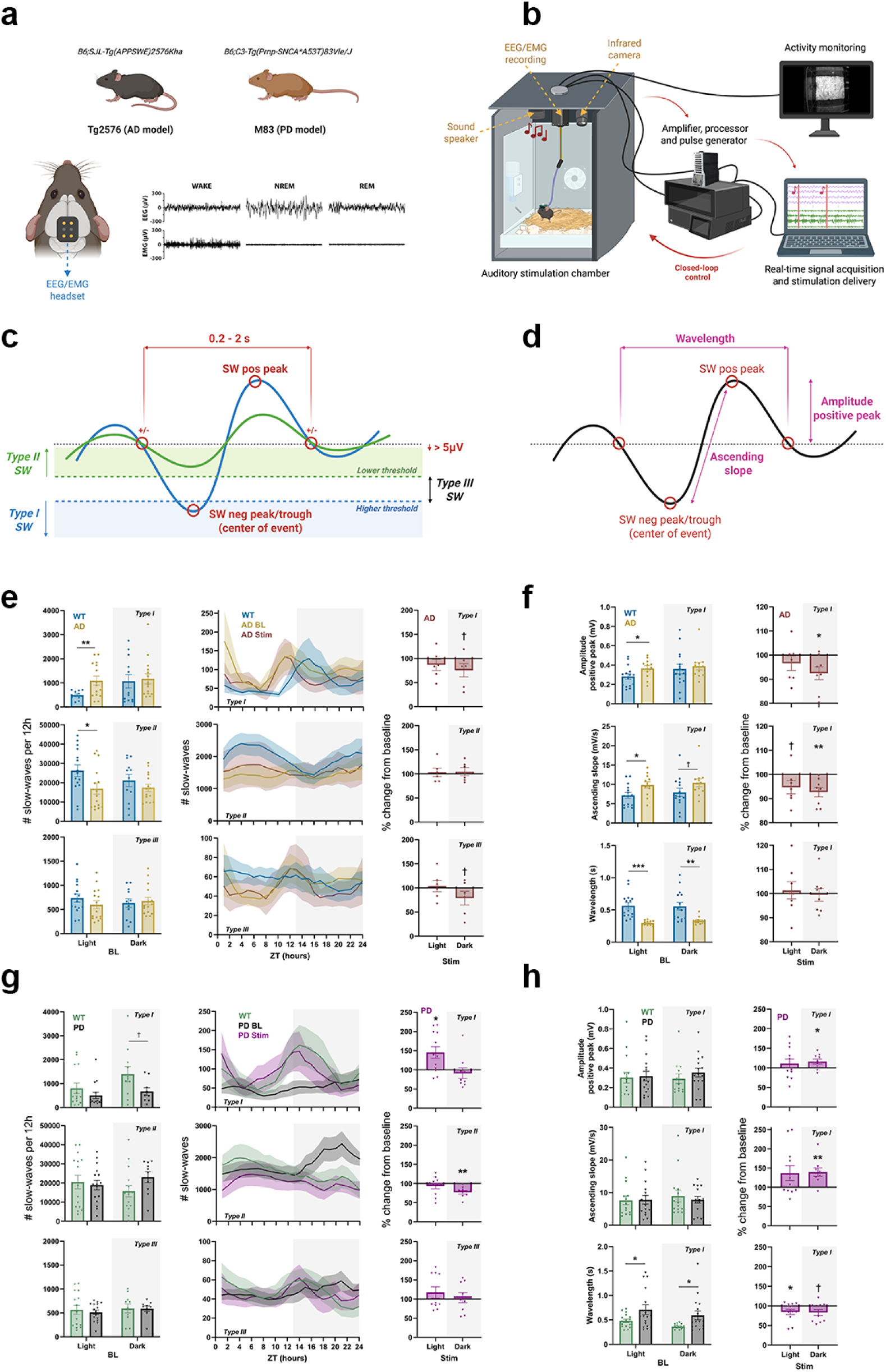
Slow-wave modulation by mCLAS in AD and PD mice. **a)** Top, mouse models of AD (Tg2576, n = 7 mice) and PD (M83, n = 7 mice) used in electrophysiological recordings (WT counterparts bred in each strain, n = 7 per model). Bottom, EEG/EMG headset location and sample recordings demonstrating vigilance state characterization; **b)** Illustration of the mouse closed-loop auditory stimulation (mCLAS) setup, combining simultaneous activity monitoring, EEG/EMG recordings, and real-time signal acquisition for auditory stimulation delivery; **c)** Schematic detailing slow-wave (SW) characterization in type I, II or III; **d)** SW characteristics derived from categorization in **c**; **e)** Left panels show the number of SWs of the corresponding type (I, II or III) at baseline (BL), for WT and AD mice in the light and dark periods. Middle panels depict the 24h dynamics of the number of type I-III SWs of WT, AD mice at baseline (AD BL) and during stimulation days (AD Stim). Right panels represent the number of type I-III SWs as percentage of change from BL during light and dark periods in AD mice (Stim); **f)** Left panels show the positive peak amplitude, ascending slope or wavelength of type I SWs at BL for WT and AD mice. Right panels represent the same measures as percentage of change from BL during light and dark periods in AD mice (Stim); **g)** as in **e** but for PD mice; **h)** as in **f** but for PD mice. In the bar plots, each scatter dot represents average daily values of each animal, each bar the mean, and the whiskers the SEM. In the 24h dynamics plots, shaded areas correspond to the SEM and the grey shaded panel corresponds to the dark period. ***p<0.001, **p<0.01, *p<0.05, †<0.2. Illustrations in a and b, and schematics in c and d were created with BioRender (https://BioRender.com).

In AD mice, we found that the average number of type I SWs was significantly higher than in WT controls at baseline during the light period (**p<0.01, two-sample t-test, **Figure 1e left**), which tended to decrease upon mCLAS in the dark period (% of change from baseline, †p=0.152, one-sample t-test, **Figure 1e right**). Conversely, type II SWs were significantly lower in AD mice than in WT controls at baseline during the light period (*p<0.5, **Figure 1e left**), with no significant change exerted by mCLAS (**Figure 1e right**). Type III SWs did not differ significantly at baseline between WT and AD mice (**Figure 1e left**), but mCLAS exerted a trend decrease during the dark period (†p=0.132, **Figure 1e right**). Inspection of 24h dynamics for each SW type support these results (**Figure 1e middle**). Regarding type I SW characteristics, positive peak amplitude and ascending slopes were significantly higher at baseline during the light period in AD mice than in WT controls (*p<0.05, **Figure 1f left**), with mCLAS facilitating a significant decrease in these metrics during the dark period (amplitude of positive peak: *p<0.05; ascending slope: **p<0.01, **Figure 1f right**). Baseline positive peak amplitude and ascending slope of type II and III SWs were similarly elevated in the light and dark periods (**p<0.01 or *p<0.05, see **Supplementary Figures 1a, b left**), with no significant changes upon mCLAS (**Supplementary Figures 1a, b right**). Additionally, wavelength values of type I SWs in AD mice were significantly lower (i.e. faster waves) at baseline compared to WT controls (light: ***p<0.001; dark: **p<0.01, **Figure 1f left**), with no significant change exerted by mCLAS (**Figure 1f right**). Wavelengths of type II and III SWs showed similar patterns (**Supplementary Figure 1c**).

Baseline and mCLAS-mediated SW dynamics were quite distinct in PD mice. Average number of type I SWs at baseline trended to be lower during the light and dark periods in PD mice compared to WT controls (†p=0.07, two-sample t-test, **Figure 1g left**), with mCLAS promoting an increase during the light period (%change from baseline, *p<0.05, one-sample t-test, **Figure 1g right**). Analysis of type I SW dynamics per 24h show that mCLAS restores SW patterns in PD mice towards WT levels, rescuing a baseline pathological flat temporal distribution of events throughout the light and dark periods (**Figure 1g middle**). Conversely, the number of type II SWs appeared non-significantly higher in PD mice at baseline during the dark period compared to WT controls (**Figure 1g left**), with a significant decrease upon mCLAS in the dark period (**p<0.01, **Figure 1g right**). Moreover, analyses of 24h dynamics show that mCLAS restores the number of type II SWs in PD mice towards WT levels in the dark period (**Figure 1g middle**). Type III SWs did not differ significantly at baseline between WT and PD mice, with mCLAS exerting no significant modulation (**Figure 1g bottom**). Regarding type I SW characteristics, baseline positive peak amplitude and ascending slopes did not differ in PD mice compared to WT controls (**Figure 1h left**), with mCLAS still promoting a significant increase in these metrics during the dark period (amplitude of positive peak: *p<0.05; ascending slope: **p<0.01, **Figure 1h right**). Similarly, positive peak amplitude and ascending slope of type II and III SWs were not altered at baseline (**Supplementary Figures 1d, e left**), with significant increases upon mCLAS (***p<0.001 or *p<0.05, see **Supplementary Figures 1d, e right**). Additionally, wavelength of type I SWs was significantly higher in PD mice (i.e. slower waves) in relation to WT controls at baseline during light and dark periods (light, dark: *p<0.05, **Figure 1h left**), with mCLAS decreasing their length (i.e. inducing faster waves) in both light and dark periods (light: *p<0.05; dark: †p=0.057, **Figure 1h right**). Wavelengths of type II and III SWs showed similar patterns (**Supplementary Figure 1f**).

### Stimulation alters sleep spindle density irrespective of baseline activity

To assess baseline sleep spindle characteristics in AD and PD mice and the potential modulatory effects of mCLAS on their dynamics, we characterized all (10-15 Hz), slow (10-12 Hz) and fast (12-15 Hz) spindles (**Figure 2a**, see Methods), and calculated the absolute number of spindles, maximum peak amplitude and spindle duration within each frequency band (**Figure 2b**), across experimental groups, protocol days and light/dark periods.

**Figure 2.**
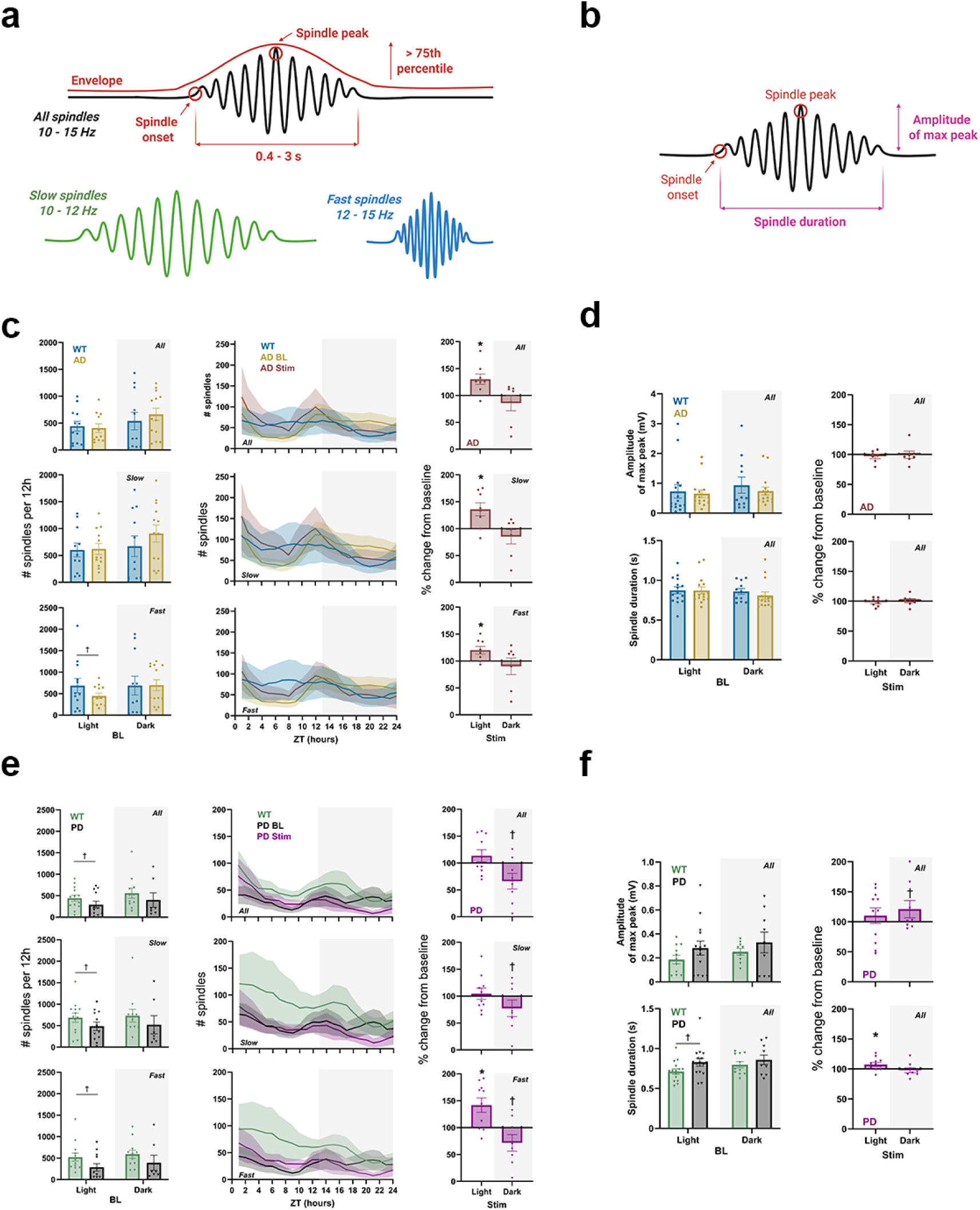
Sleep spindle modulation by mCLAS in AD and PD mice. **a)** Schematic detailing spindle characterization in all, slow or fast spindles; **b)** Spindle characteristics derived from categorization in **a**; **c)** Left panels show the number of spindles of the corresponding type (all, slow or fast) at baseline (BL), for WT and AD mice in the light and dark periods. Middle panels depict the 24h dynamics of the number of spindle types for WT, AD mice at baseline (AD BL) and during stimulation days (AD Stim). Right panels represent the number of spindle types as percentage of change from BL during light and dark periods in AD mice (Stim); **d)** Left panels show the amplitude of the max peak and spindle duration of all spindles at BL for WT and AD mice. Right panels represent the same measures as percentage of change from BL during light and dark periods in AD mice (Stim); **e)** as in **c** but for PD mice; **f)** as in **d** but for PD mice. In the bar plots, each scatter dot represents average daily values of each animal, each bar the mean, and the whiskers the SEM. In the 24h dynamics plots, shaded areas correspond to the SEM and the grey shaded panel corresponds to the dark period. *p<0.05, †<0.2. Schematics in a and b were created with BioRender (https://BioRender.com).

In AD mice, the average number of all, slow and fast spindles did not differ significantly compared to WT controls at baseline during both light and dark periods (**Figure 2c left**), with only fast spindles showing a trend decrease in the light period (†p=0.187, two-sample t-test, **Figure 2c left).** Despite no baseline alterations in AD mice, mCLAS elicited a potentiation in all, slow and fast spindle events in the light period (% change from baseline, all, slow, fast: *p<0.05, one-sample t-test, **Figure 2c right**). These results are confirmed by inspection of 24h dynamics for all spindle types (**Figure 2c middle**). Amplitude of maximum spindle peak and spindle duration did not differ at baseline nor upon mCLAS across all (**Figure 2d**), slow and fast spindles (**Supplementary Figures 2a, b**).

Similarly, in PD mice we found that the average number of all, slow and fast spindles did not differ significantly compared to WT controls at baseline during light and dark periods, although we observed trend decreases in the light period for all, slow and fast spindles (all:†p=0.176; slow:†p=0.178; fast:†p=0.07; two-sample t-test, **Figure 2e left**). In spite of no observable baseline alterations, mCLAS led to a significant increase in fast spindles in the light period of PD mice (% change from baseline, *p<0.05, one-sample t-test, **Figure 2e right**), and a trend reduction in all, slow and fast spindles in the dark period (all:†p=0.051; slow:†p=0.179; fast:†p=0.109, **Figure 2e right**). Inspection of 24h dynamics for each spindle band support these results (**Figure 2e middle**). Amplitude of maximum spindle peak and spindle duration did not differ significantly between PD mice and WT controls at baseline for all (**Figure 2f left**) and slow spindles (**Supplementary Figures 2c, d left**), whereas fast spindle duration was significantly increased in the light period in PD mice (*p<0.05, **Supplementary Figure 2d left**). Moreover, stimulation exerted an increase in all spindle duration in PD mice (% change from baseline, *p<0.05, **Figure 2f right**), with no significant effects observed in slow and fast spindles (**Supplementary Figures 2c, d right)**.

### mCLAS shapes SW-spindle coupling strength, directionality and event-locked spindle power

To assess SW-spindle coupling in AD and PD mice and the potential modulatory effects of mCLAS on their dynamics, we defined SW-spindle coupling (**Figure 3a left,** see Methods) between SW types I, II and III, and all, slow and fast spindles using criteria previously described^45^. As we observed negligible baseline alterations and mCLAS modulation on type III SWs, as well as similar baseline and modulation dynamics across all, slow and fast spindles, we only report coupling metrics for type I SWs – all spindles (**Figure 3**) and type II SWs – all spindles (**Supplementary Figure 3**) in both transgenic lines. We assessed the number of spindles coupling at the SW peak, cross-frequency coupling strength via the modulation index (MI), and directionality through the phase slope index (PSI) (**Figure 3a middle** see Methods). Additionally, we performed event-locked spectral analysis by computing time frequency representations (TFRs) from which we extracted spindle power at SW trough, peak and at maximum spindle power (**Figure 3a right** see Methods), across experimental groups, protocol days and light/dark periods.

**Figure 3.**
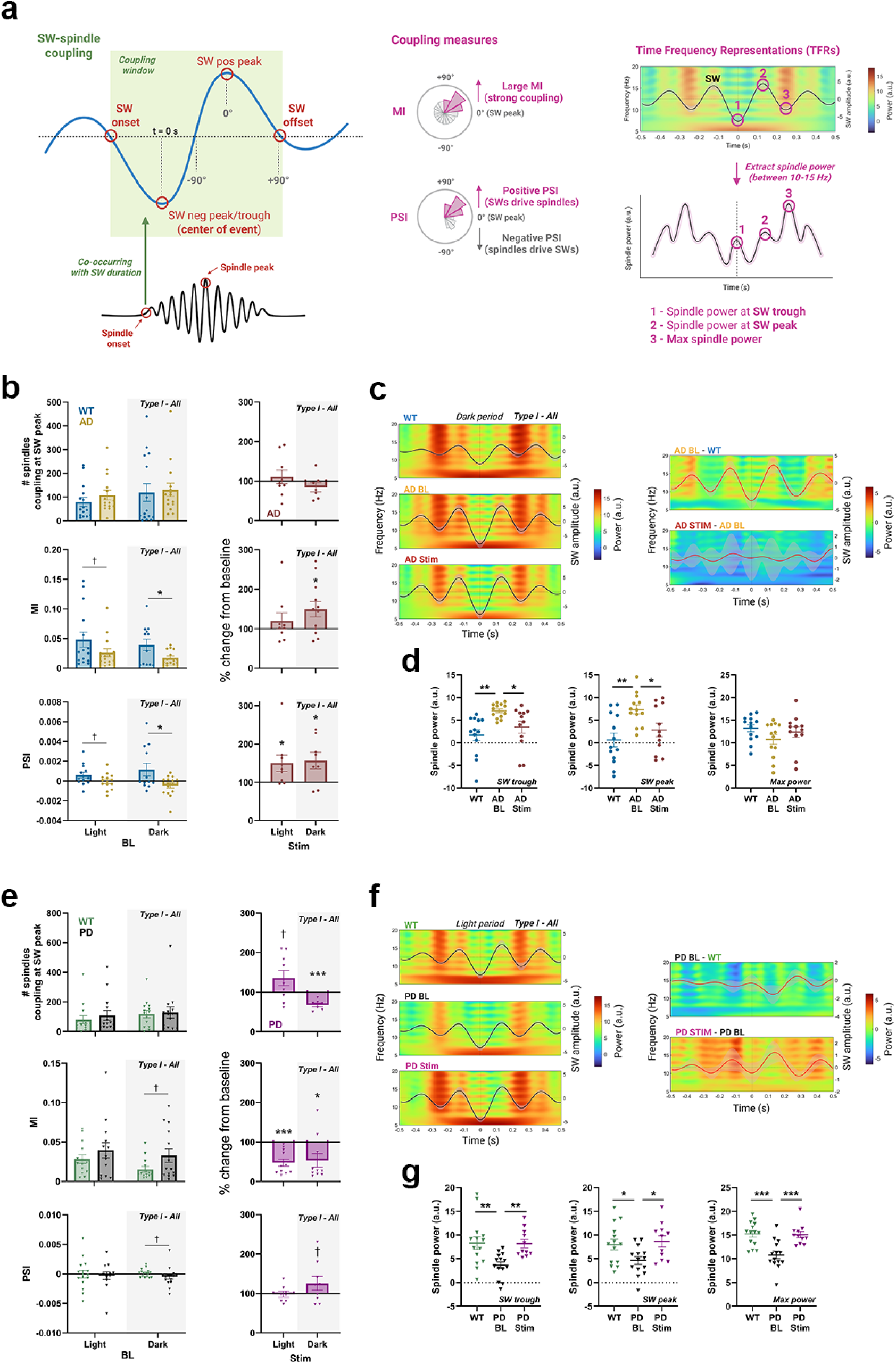
SW-spindle modulation by mCLAS in AD and PD mice. **a)** Left panel: Schematic detailing slow-wave (SW)-spindle coupling characterization; Middle and right panels: SW-coupling measures derived from categorization, including the modulation index (MI), the phase slope index (PSI), time frequency representations (TFRs) and spindle power at SW trough, peak and maximum of spindle power; **b)** Left panels show either the number of spindles coupling at SW peak (top), MI (middle) or PSI (bottom), for SWs type I – all spindles coupling at baseline (BL) in WT and AD mice in the light and dark periods; Right panels show the percentage of change from BL during light and dark periods in AD mice (Stim) across those measures; **c)** Left panels show TFRs and average SWs traces for type I – all spindles coupling during the dark period in WT, AD mice at baseline (AD BL) and during stimulation days (AD stim). Right panels show difference TFRs and average SW traces between AD BL-WT and AD Stim-AD BL; **d)** Spindle power at SW trough, SW peak and maximum spindle power extracted from TFRs in c for WT, AD BL and AD Stim; **e)** as in **b** but for PD mice; **f)** as in **c** but for PD mice; **g)** as in **d** but for PD mice. In the bar and scatter plots, each scatter dot represents average daily values of each animal, each bar the mean, the whiskers the SEM and the grey shaded panel corresponds to the dark period. In the TFR plots, black solid lines correspond to the average SW, red solid lines correspond to the average SW difference, shaded areas correspond to the SEM and the dotted line at t = 0 seconds correspond to the SW trough (event center). ***p<0.001, **p<0.01, *p<0.05, †<0.2. Schematics in a and b were created with BioRender (https://BioRender.com). TFRs in c and f were created in MATLAB (R2023b).

For type I SWs - all spindles coupling in AD mice, the average number of spindles coupling at the SW peak did not differ from WT controls at baseline, with no observable effects of mCLAS (**Figure 3b top**). Cross-frequency coupling strength as determined by MI was significantly decreased at baseline in AD mice compared to WT controls mainly in the dark period (light: †p=0.133; dark: *p<0.05, two-sample t-test, **Figure 3b middle left**), which significantly increased upon mCLAS (% change from baseline, *p<0.05, one-sample t-test, **Figure 3b middle right**). Coupling directionality determined by PSI followed the same pattern, with negative values (i.e. spindles driving SWs) in AD mice at baseline, mainly in the dark period (light: †p=0.133; dark: *p<0.05, **Figure 3b bottom left**), which were significantly increased by mCLAS (light and dark: *p<0.05, **Figure 3b bottom right**). As mCLAS resulted in a significant MI increase during the dark period only, we followed up with event-locked spectral analysis for this period. TFRs showed noticeable differences between AD mice and WT controls at baseline (**Figure 3c**), with elevated spindle power at SW trough and peak (trough and peak: **p<0.01, one-way ANOVA with Holm-Šidák post hoc test, **Figure 3d left and middle**), which were significantly decreased by mCLAS (trough and peak: *p<0.05, **Figure 3d left and middle**). At maximum spindle power, AD mice showed non-significantly lower values than WT controls at baseline (**Figure 3c, d right**), which were non-significantly potentiated by mCLAS (**Figure 3d right**).

We observed similar patters for type II SWs - all spindles coupling in AD mice. The number of spindles coupling at the SW peak was increased in AD mice compared to WT controls at baseline (*p<0.05, two-sample t-test, **Supplementary Figure 3a top left**), with no significant effects of mCLAS. MI trended to be lower in AD mice than in WT controls at baseline during the dark period (dark: †p=0.064, **Supplementary Figure 3a middle left**), which was significantly counteracted by mCLAS (% change from baseline, *p<0.05, one-sample t-test, **Supplementary Figure 3a middle right**). PSI showed non-significant negative values in the dark period at baseline for AD mice, which were significantly increased upon mCLAS (light: **p<0.01 dark: *p<0.05, **Supplementary Figure 3a bottom right**). As mCLAS resulted in a significant MI increase during the dark period, we followed up with event-locked spectral analysis. TFRs showed differences between AD mice and WT controls at baseline (**Supplementary Figure 3b**), with elevated spindle power at SW trough, peak and maximum spindle power (trough: ****p<0.0001; peak: ***p<0.001; max spindle power: *p<0.05, one-way ANOVA with Holm-Šidák post hoc test, **Supplementary Figure 3c**) which were significantly decreased by mCLAS (trough: **p<0.01; peak: *p<0.05; max spindle power: *p<0.05, **Supplementary Figure 3c**).

In PD mice, SW-spindle coupling metrics were distinctly different from those of AD. For type I SWs - all spindles, the average number of spindles coupling at the SW peak did not differ from WT controls at baseline, but mCLAS potentiated a significant decrease in the dark period (% change from baseline, ***p<0.001, two-sample t-test, **Figure 3e top**). MI trended to be higher in PD mice than in WT controls at baseline during the dark period (dark: †p=0.068, two-sample t-test, **Figure 3e middle left**), which significantly decreased upon mCLAS (% change from baseline, light: ***p<0.001; dark: *p<0.05, one-sample t-test, **Figure 3e middle right**). PSI showed trend negative values for PD mice at baseline in the dark period (†p=0.198, **Figure 3e bottom left**), which were trend increased by mCLAS (†p=0.187, **Figure 3e bottom right**). As mCLAS resulted in a strong MI decrease during the light period, we followed up with event-locked spectral analysis in this period. TFRs showed striking differences at baseline between PD mice and WT controls (**Figure 3f**), with decreased spindle power at SW trough, peak and maximum spindle power (trough: **p<0.01; peak: *p<0.05; max spindle power: ***p<0.001, one-way ANOVA with Holm-Šidák post hoc test, **Figure 3g**), changes that were significantly counteracted by mCLAS (trough: **p<0.01; peak: *p<0.05; max spindle power: ***p<0.001, **Figure 3g**).

For type II SWs - all spindles coupling in PD mice, we observed similar patterns. The number of spindles coupling at the SW peak was trend decreased in PD mice in relation to WT controls at baseline (light: †p=0.068 ; dark: †p=0.123, two-way sample t-test, **Supplementary Figure 3d top left**), with a non-significant recovery exerted by mCLAS in the dark period (**Supplementary Figure 3d top right**). MI trended to be higher in PD mice than in WT controls at baseline (light: †p=0.117;dark: †p=0.158, **Supplementary Figure 3d middle left**), which decreased upon mCLAS in the dark period (% change from baseline, †p=0.086, one-sample t-test, **Supplementary Figure 3d middle right**). PSI showed non-significant negative values at baseline in the dark period for PD mice (†p=0.175, **Supplementary Figure 3d bottom left**), with this metric significantly increasing by mCLAS throughout the day (light: *p<0.05 dark: **p<0.01, **Supplementary Figure 3d bottom right**). As mCLAS resulted in a MI decrease during the dark period, we followed up with event-locked spectral analysis. TFRs showed differences at baseline between PD mice and WT controls (**Supplementary Figure 3e**), with lower spindle power at SW trough, peak and maximum spindle power (trough: **p<0.01; peak: ****p<0.0001; max spindle power: **p<0.01, one-way ANOVA with Holm-Šidák post hoc test, **Supplementary Figure 3f**), which was significantly counteracted by mCLAS (trough: *p<0.01; peak: **p<0.01; max spindle power: *p<0.05, **Supplementary Figure 3f**).

### Stimulation differentially drives sleep pressure dissipation

To assess alterations in sleep pressure dissipation dynamics in AD and PD mice and the potential modulatory effects of mCLAS, we assessed SWE across experimental groups.

In AD mice, SWE across 24h significantly differed between WT controls and AD mice at baseline and upon mCLAS (****p<0.0001, two-way mixed ANOVA with Greenhouse-Geisser correction, **Figure 4a left**). At ZT24, we observed that cumulative SWE was decreased in AD mice when compared to WT controls (****p<0.0001, two-way ANOVA with Holm-Šidák post hoc test, **Figure 4a right**), which was significantly counteracted upon mCLAS (*p<0.05, **Figure 4a right**).

**Figure 4.**
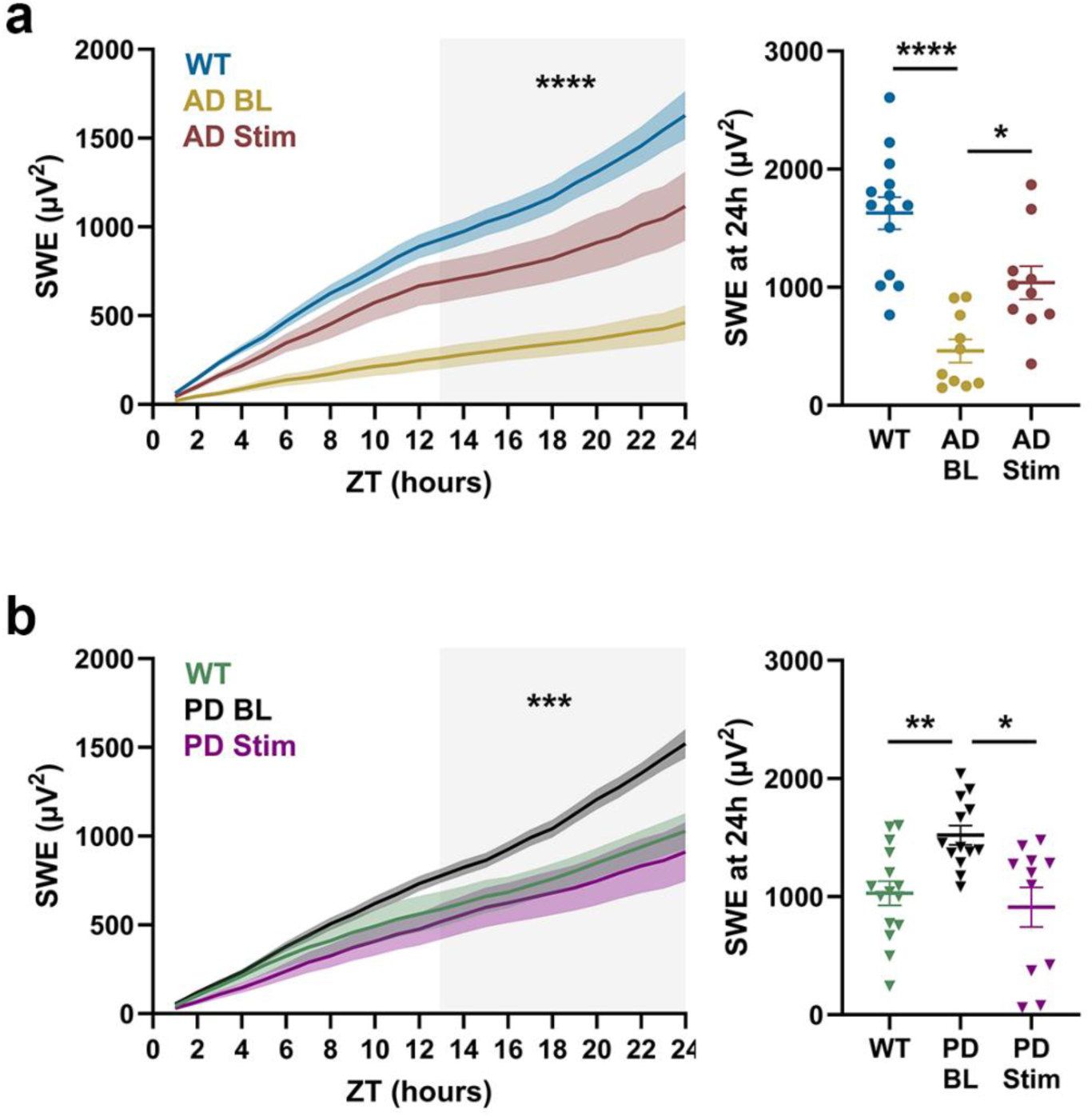
mCLAS drives sleep homeostasis in AD and PD. **a)** Left panel depicts the cumulative values of SWE across 24h for WT, AD mice at baseline (AD BL) and during stimulation days (AD Stim). Right panel shows SWE at hour 24 for WT, AD BL and AD Stim; **b)** as in a but for PD mice. In the 24h plots, shaded areas correspond to the SEM and the grey shaded panel corresponds to the dark period. In the scatter plots, each dot represents average daily values of each animal, each bar the mean, and the whiskers the SEM. ****p<0.0001; ***p<0.001, **p<0.01, *p<0.05.

In PD mice, SWE throughout 24h significantly differed across WT controls and PD animals at baseline and upon mCLAS (***p<0.001, two-way mixed ANOVA with Greenhouse-Geisser correction, **Figure 4b left**). Contrarily to what we observed in AD, at ZT24 SWE was increased in PD mice in relation to WT controls (**p<0.01, two-way ANOVA with Holm-Šidák post hoc test, **Figure 4b right**), which was significantly normalised by mCLAS (*p<0.05, **Figure 4b right**).

## Discussion

In this study, we demonstrate that up-phase-targeted mCLAS rescues key pathological alterations of SWs, spindles, cross-frequency coupling and sleep dissipation in two mouse models of neurodegeneration. To achieve this, we applied mCLAS parameters previously optimised^40^ for each disease model in a 2-day baseline/stimulation design, and conducted an in-depth analysis of the acute effects of mCLAS on relevant metrics, previously described as impaired in aging and disease.

In WT, AD and PD mice, we verified that type I (global, subcortico-cortical synchronization) and type II SWs (local, cortico-cortical synchronization) ^19–21^ co-occur during the light and dark periods. In WT mice, we observed that type II SWs decrease in number during the light (rest) period, reflecting sleep pressure dissipation and homeostatic decline ^20^. In contrast, type I SWs lack this regulation, peaking at at the beginning of the dark (active) period, perhaps reflecting increased activity of the subcortical arousal system and/or decreased number and amplitude of type II SWs limiting type I SW spread^20^.

In AD mice, we observed that type I SWs were elevated at baseline relative to their WT littermates, indicating reduced sleep depth and heightened interaction between arousal structures and the cortex^20^. This aligns with previous findings of increased amplitude peaks and steeper ascending slopes in older healthy adults^46^ and in AD mice^10^, as well as elevated wakefulness in our murine AD cohort^10,40^, factors linked to higher amplitudes, steeper SW slopes and increased sleep pressure^47,48^. Conversely, type II SWs were reduced at baseline, reflecting impaired sleep maintenance and environmental disconnection^20^. This finding mirrors reports of reduced SW density in older adults^16^ and AD mice^10^, likely driven by type II SW prevalence^20^. Our mCLAS paradigm acutely reduced the number of type I SWs, enhancing sleep depth and continuity, as evidenced by previously reported decreased NREM sleep fragmentation, microarousal events and vigilance state transitions^40,49^. mCLAS also attenuated abnormal high amplitudes and steep ascending slopes of type I SWs, reducing network recruitment and impairing synchronous propagation^20^. While mCLAS did not increase type II SW counts, it potentiated their amplitude and slope, suggesting enhanced local network recruitment and compensatory cortical synchrony^20^. This reorganization altered the 24-hour temporal pattern of type II SWs, mildly counteracting the arrhythmic baseline pattern.

In stark contrast, PD mice displayed reduced type I SWs at baseline, indicative of pathological deep sleep and excessive environmental disconnection, as reported in PD patients and mice^49,50^. mCLAS increased type I SW incidence in PD mice and restored its 24-hour rhythmicity to WT levels, suggesting an enhanced interaction between arousal structures and the cortex^20^. This aligns with our earlier results showing increased vigilance state transitions and decreased sleep continuity, alongside heightened arousal following stimulation^40,49^. mCLAS also normalized the abnormally long wavelengths of type I and II SWs in PD mice, inducing faster oscillations, and increased their amplitude and slope, suggesting altered excitation/inhibition (E/I) balance during light-to-dark transitions and the dark period^20^. This aligns with reports linking excessive daytime sleepiness in PD patients to reduced homeostatic decline of SWA and spindle frequency^51^, potentially due to excessive type II SW prevalence during the active period. Remarkably, mCLAS decreased the number of type II SWs in PD mice, reducing environmental disconnection^20^, and normalized their 24-hour temporal distribution toward WT levels, especially during the dark period. Finally, the effects of mCLAS on type III SWs mirrored those on type I, reflecting a systemic shift toward global synchronization by potentiating type II-to-type I SW “conversion”.

Previous human studies in healthy young individuals have shown that CLAS can modulate SW types, with up-phase stimulation of type II SWs promoting greater cortical involvement^52^ and triggering both type I and type II SWs^53^. While^53^ hypothesized that evoking type I SWs might benefit patients with neurodegeneration, our results in prodromal AD mice challenge this idea: baseline increases in type I SWs were associated with sleep fragmentation and heightened arousal^40,49^, suggesting that promoting type II SWs to enhance sleep depth may be more beneficial in the context of AD. Conversely, potentiating type I SWs to increase vigilance transitions could be advantageous for PD patients^49^. These disease-specific responses underscore that mCLAS effects in healthy individuals may not translate directly to pathological contexts, highlighting the need for long-term studies coupling sleep, cognition and protein burden analyses.

In contrast to the pronounced effects of mCLAS on SW dynamics, its impact on sleep spindles in both AD and PD mice was more nuanced. Despite the absence of significant baseline impairments in spindle density, with only a trend towards reduction in the light period in PD mice, mCLAS consistently potentiated spindle activity during the light period for both AD and PD, while inducing a non-significant decrease during the dark period, indicating a marked temporal reorganization. This modulation occurred without altering spindle amplitude or duration, suggesting that mCLAS selectively redistributes spindle density rather than modifying their intrinsic characteristics. These findings align with the hierarchical relationship between SWs and spindles, where SWs are known to drive spindle generation in healthy individuals^9,54^. Given that both AD and PD mice in this study were at a prodromal disease stage, it is plausible that early pathological processes first manifest in SW alterations, with downstream effects on spindles emerging only at later stages^17,55,56^. Thus, the potentiation of spindles by mCLAS, particularly during the light period, may reflect a compensatory mechanism to restore SW-spindle coupling before overt spindle deficits arise.

Accordingly, our results reveal that mCLAS differentially restores impaired SW-spindle coupling in AD and PD mice, acting in direct opposition to their baseline deficiencies. At baseline, AD mice exhibited reduced cross-frequency coupling strength during the dark period, mirroring impairments observed in human MCI^55^. Strikingly, mCLAS increased MI particularly during the dark period, effectively restoring coupling strength. This restoration was paralleled by a reversal of altered coupling directionality: negative PSI values at baseline, indicating spindles driving SWs, as seen in aging^18^, shifted toward positive values under stimulation, reinstating the canonical hierarchy where SWs drive spindles^9^. Event-locked spectral analyses further revealed an mCLAS-mediated renormalization of spindle power dynamics, reducing the abnormally high spindle power at SW trough and peak. These effects were consistent for both type I/II SWs – all spindles coupling, underscoring a systematic recalibration of coupling orchestration rather than isolated changes in individual oscillatory features. Notably, the number of spindles coupling at the SW peak remained unchanged by mCLAS, suggesting that the intervention primarily modulates the quality (strength/directionality) rather than the quantity of coupling events. This aligns with the idea that the transition from young, to healthy aging to prodromal neurodegeneration stages is characterized by disrupted coordination between SWs and spindles, rather than a loss of spindles themselves^18,55^. By restoring coupling strength and directionality, mCLAS may thus counteract the early hierarchical breakdown in SW-spindle interactions^18^.

In stark contrast to AD, PD mice exhibited abnormally elevated baseline coupling strength, a pattern suggesting excessive yet inefficient SW-spindle orchestration. Here, mCLAS exerted the opposite effect, decreasing coupling strength during both light and dark periods in type I SW – all spindle coupling. Baseline PSI values in PD mice trended negative (spindles driving SWs), but mCLAS shifted directionality toward positive values, mirroring the restoration of SW-driven hierarchy also exerted in AD. However, the mechanism differs fundamentally: in PD, mCLAS reduced excessive coupling strength while enhancing spindle power at SW trough, peak and maximum spindle power, suggesting that the intervention mitigates pathological hypercoupling rather than weakness. The significant decrease prompted by mCLAS in the number of spindles coupling at the SW peak during the dark period further supports this interpretation, indicating that mCLAS may prune excessive non-physiologic coupling events to restore a more healthier and balanced SW-spindle dialogue.

The dichotomous effects of mCLAS in AD and PD mice highlight its capacity to adaptively normalise coupling based on baseline deficiencies. In AD, where coupling is weak and spindle-driven, mCLAS strengthens it to restore SW-driven hierarchy. In PD, where coupling is overly strong but inefficient, mCLAS reduces its strength while enhancing spindle power, rebalancing the system. Both disease models share a reversal of pathological coupling directionality, reinforcing that mCLAS targets the hierarchical organization of SW-spindle interactions, a process disrupted in human ageing^9,18^. Critically, these effects were consistent across the cross-frequency coupling types analysed (type I/type II – all spindles), suggesting that mCLAS acts on fundamental coupling mechanisms rather than specific oscillatory subtypes. The restoration of coupling strength, directionality and spindle power suggests that mCLAS acts equally on cross-frequency strength and coordination, mechanisms proposed to underpin memory consolidation^9,54,55^. By demonstrating that mCLAS can reverse pathological coupling hierarchies in prodromal neurodegeneration, we provide proof-of-concept that up-phase targeted auditory stimulation may counteract not only individual oscillatory deficits but also their dysfunctional orchestration, a critical step toward restoring sleep’s role in memory and brain health.

Building on our previous work showing that short-term mCLAS enhances SWA in both AD and PD models^40^, we here demonstrate that mCLAS also bidirectionally modulates SWE, counteracting baseline homeostatic impairments. In AD mice, where baseline SWE was significantly reduced compared to WT controls, mCLAS increased cumulative SWE over 24 hours, bringing it closer to physiological levels. This increase in SWE aligns with the restoration of coupling strength observed, suggesting that mCLAS mitigates the deficient homeostatic decline characteristic of prodromal AD, potentially counteracting the sleep fragmentation and reduced SWA associated with amyloid pathology^10,25^. Conversely, in PD mice, where baseline SWE was abnormally elevated relative to WT controls, mCLAS significantly decreased SWE, fully renormalizing sleep pressure dissipation. This reduction mirrors the attenuation of excessive coupling strength and restoration of spindle power observed in PD, indicating that mCLAS corrects abnormal sleep homeostasis. Together, these findings demonstrate that mCLAS does not merely enhance SWA universally but selectively modulates sleep homeostasis in a context-dependent disease-specific manner, offering a targeted approach to restore physiological sleep dynamics in neurodegeneration.

Accordingly, the current results hint two potential mechanisms through which mCLAS modulates sleep homeostasis in AD and PD mice: 1) a direct influence on cortical generators^20,41,52,57^, leading to type II SW potentiation and engagement of cortical networks, which hamper subsequent propagation and synchronization of type I SWs, or at least decrease network recruitment to initiate these phenomena^20^; or/and 2) a direct effect on arousal promoting structures potentiating type I SWs, which could consequently still trigger increased synchrony at a cortical level^20,57,58^. Moreover, these two mechanisms may not to be mutually exclusive, interchanging throughout 24h. Based on our results and current literature, we propose the following flow of events upon mCLAS in:

- **AD mice**: stronger background activity of type I SWs, stimuli have a direct effect on cortical networks (**Figure 5a**), implying that:

1. The SWs mainly potentiated by mCLAS in the light period are type II, corresponding to local cortical synchrony processes^9,18,20,29,59^;
2. When thalamic spindle activity is high^60^, with increased SW-spindle coupling strength and SW-driven hierarchy, downstream locus coeruleus (LC) tonic firing is low, with low paraventricular thalamic activation impeding arousal^61,62^;
3. Arousal and spindle power at SW trough and peak decrease. The cortex is less responsive to arousing stimuli, resulting in decreased cortical microarousals, NREM sleep fragmentation and impeded vigilance state transitions, linked with decreased occurrence, amplitude and synchrony of type I SWs (global synchrony processes promoting arousal are impeded, as networks are already engaged and less recruited)^20^;
4. NREM sleep duration and continuity are promoted, leading to synaptic strength regulation^20,63,64^. This effect allows for sleep restoration and homeostatic regulation, with more efficient sleep pressure dissipation throughout NREM sleep^22,41^;
5. This cycle of points 1-4 continues until sleep pressure is dissipated, with type I SWs increasing towards the end of the light period, implying increased arousal and transitions into WAKE and REM sleep, which extend throughout the dark period. The circadian rhythmicity of vigilance states is restored.
- **PD mice**: stronger background activity of type II SWs, stimuli have a direct effect on arousal promoting structures (**Figure 5b**), implying that:

1. mCLAS potentiates increased activity of arousal promoting areas (as the LC), particularly in the light-dark period transition and during the dark period, signalling higher levels of norepinephrine release. When LC tonic firing is high, thalamic spindle activity and SW-spindle coupling strength are low^60^, with high paraventricular thalamic activation promoting arousal^61,62^;
2. Arousal promoting areas project to the orbitofrontal cortex (OFC), where type I SWs are mainly generated^58^, resulting in increased spindle power at trough and peak with SW-driven coupling hierarchy. Global synchrony processes promoting arousal are induced, with more networks recruited to engage in these events, which propagate throughout the cortex and subsequently initiate local processes;
3. As networks are already engaged in large scale global SW synchronization, local type II SWs are still induced but in decreased amount, particularly during the dark period^20^. The cortex is more responsive to arousing stimuli, resulting in increased vigilance state transitions, which could relate to restored dopaminergic neurotransmission upon stimulation^50,65–67^. Moreover, disruption of overly consolidated NREM sleep and synaptic strength regulation are also promoted^20,63,64^;
4. There is a decrease in sleep depth during the light-to-dark transition and the dark period, implying increased arousal and transitions into WAKE and REM sleep. As a consequence, sleep pressure builds up during the dark period;
5. During the light period, cumulative SWE decreases due to a decrease in overly consolidated NREM sleep states, enabling physiological sleep restoration and homeostatic regulation during the beginning of the light period, with enhanced sleep pressure dissipation throughout NREM sleep and decreased sleep need in the dark period^22,41^. The circadian rhythmicity of vigilance states is restored.

**Figure 5.**
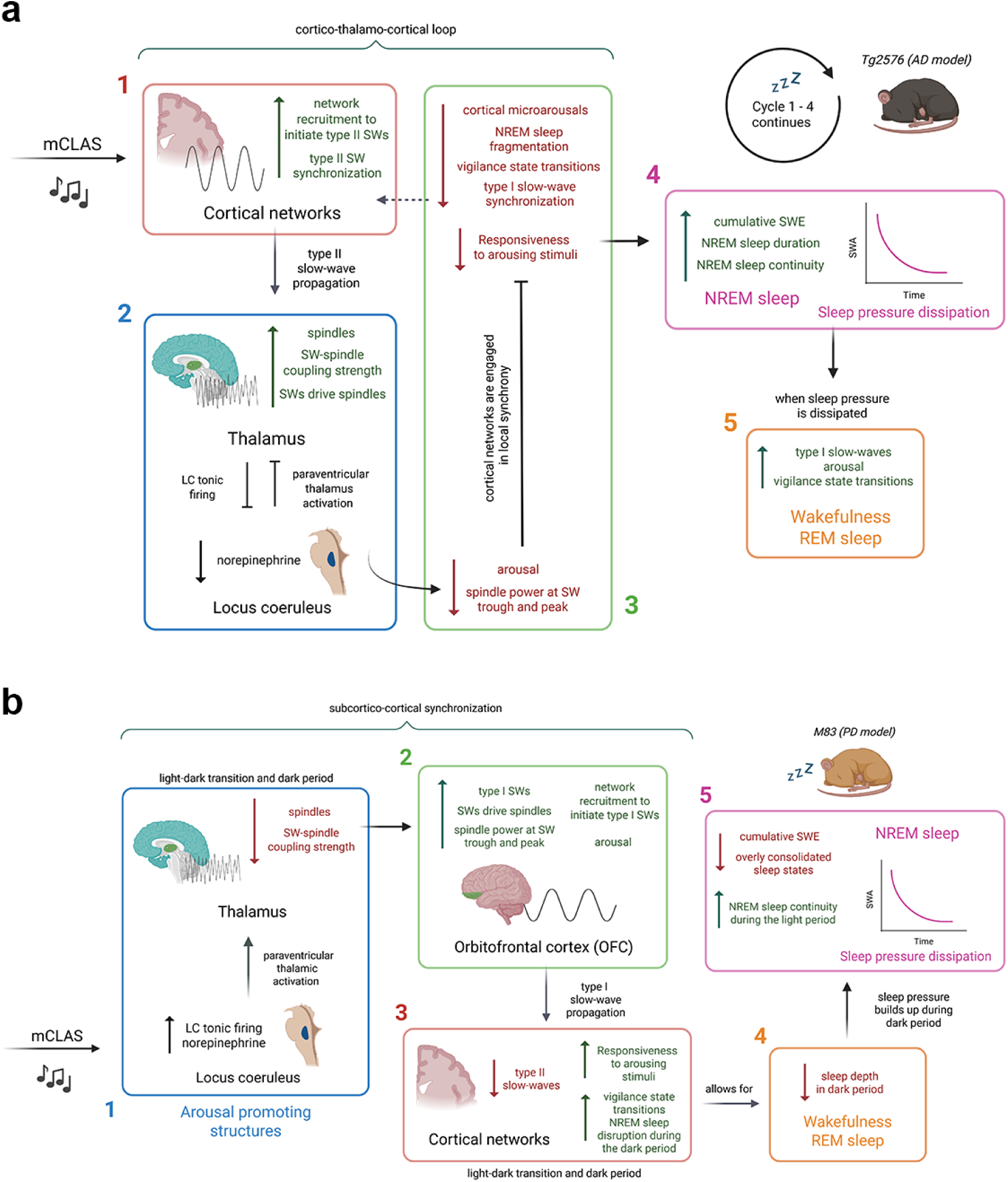
Bidirectional and context-dependent regulation of sleep homeostasis in AD and PD. **a)** Proposed sequence of events in AD mice upon mCLAS; **b)** Proposed sequence of events in PD mice upon mCLAS. Illustrations and schematics were created with BioRender (https://BioRender.com).

The SWs mainly tracked by our set-up and targeted with stimuli are most likely SWs type I, as the algorithm is tunned to phase-lock stimuli to big amplitudes^68^. These external auditory stimuli targeted to the up-phase of SWs prompt the initiation of further down- and subsequent up-states, potentiating continuity of SW cycles in NREM sleep^69^. Our results hint that the subsequent SW cycles prompted by mCLAS depend on the pre-state of each system^20^, the baseline state, which is differently impaired by disease in AD and PD mice. We suggest that the cascade of effects leading to restored sleep homeostasis in AD and PD mice may be due to mCLAS acting initially as an external context-dependent engager, pushing a diseased unbalanced system into a new equilibrium, which is dependent on its altered initial state. Further work is needed to assess the effects of improved SW dynamics, restored SW-spindle coupling and increased sleep pressure dissipation efficiency on behavioural phenotypes of AD and PD mice, as well as on histopathological and molecular hallmarks of both neurodegenerative processes, closing the loop on mCLAS’ potential neuroprotective effect in neurodegeneration.

Our SW characterization method adapted a human studies’ synchronization score metric based on scalp involvement and slope measurements^20,53,70^, to a single-electrode EEG setup. Since scalp involvement could not be assessed, we used negative peak amplitude as the primary metric for classifying SW types, as it is the most representative feature in both human and animal studies^10,20,52^. While this approach preserved the distinction between type I and II SWs, it limited our ability to directly compare type III SWs to those described in human literature, where scalp involvement is critical^70^. Our type III SWs, characterized by intermediate amplitudes, may represent transitional states between type II and type I dynamics^53^, but this interpretation requires further validation. Moreover, our analysis focused on SWs in the 0.5–4 Hz range, which may not fully dissociate the competing roles of slower (<2 Hz) and faster (2–4 Hz) sub bands in sleep-dependent processes such as memory consolidation and amyloid clearance^71,72^. However, our already complex analysis pipeline aligns with established SW characterization metrics in mouse sleep research^42,73,74^ and with similar SW characterization methods in human studies^20,52^. Additionally, we have not performed sex-based assessments in this study, with the results including a higher percentage of female mice due to male dropouts based on aggressiveness leading to implant loss. Data loss in several animals also prevented repeated measures analysis. This drawback was mainly due to software crashes, lost EEG implants, tether disconnections or poor signal quality related with prolonged recordings. Lastly, we did not implement adaptive feedback methods to estimate ongoing phase-targets in our recordings^32,75,76^, which could potentially improve applicability of mCLAS during NREM sleep.

In summary, we demonstrate that mCLAS selectively acts through two complementary pathways: modulation of cortical synchrony, by potentiating local network engagement via type II SWs while reducing pathological type I wave propagation in AD, and enhancing global synchrony in PD. Beyond individual SW dynamics, mCLAS recalibrates SW-spindle coupling, reversing pathological directionality and normalizing coupling strength, while bidirectionally regulating sleep homeostasis by enhancing SWE in AD and reducing its excess in PD. By targeting both the cortical generators of sleep oscillations and their orchestration with subcortical systems, mCLAS emerges as a precision tool to restore physiological sleep patterns, offering a foundation for future therapies aimed at counteracting sleep disruption and its downstream effects in neurodegenerative diseases.

## Methods

### Animal models and surgeries

We used the Tg2576 model of AD^43^, overexpressing a mutant form of the human amyloid precursor protein gene; and the M83 model of PD^44^, expressing the A53T-mutant variant of the human α-synuclein gene (**Figure 1a**). The study design included male and female transgenic mice of each strain (AD: n = 7, 4 females; and PD: n = 7, 6 females) at a prodromal disease stage (6.5 months-of-age at implantation, plaque- or heavy inclusion-free) and their age-matched healthy littermates (wild-type, WT; bred on AD strain: n = 7, 5 females; bred on PD strain: n = 7, 5 females). Mice were group-housed in ventilated cages under a 12/12h light/dark cycle with food and water available *ad libitum*. All experimental procedures were performed according to federal Swiss regulations under approval of the veterinary office of the Canton Zurich (license number ZH024/2021).

Mice were implanted as described before^10,40,49^. Briefly, we placed 1 stainless steel screw per hemisphere, each located 2 mm posterior to Bregma and 2 mm lateral from midline. The screws were connected to a pin header with copper wires for EEG recordings, and the structure fixed with dental cement (**Figure 1a).** For EMG recordings, we inserted two gold wires bilaterally into the neck muscles. All surgical procedures were performed under deep inhalation anaesthesia, with pre-emptive administration of anti-inflammatory and analgesic drugs. Moreover, we administered postsurgical analgesia for two days during both light and dark periods, and monitored body weight and home cage activity daily during the post surgical week and twice weekly thereafter^40,49^.

### EEG/EMG recordings, mCLAS paradigm, offline scoring and pre-processing

Following a two-week recovery period, mice were individually housed in sound-insulated chambers (**Figure 1b**). We continuously recorded EEG/EMG signals and delivered phase-locked auditory stimuli in real-time, as previously described^40,49,68^. Briefly, we conducted inter-hemispheric tethered recordings in differential mode, with the left hemisphere EEG signal referenced to the right hemisphere electrode, amounting to one EEG channel per mouse^40^. All signals were sampled at 610.35 Hz, amplified after applying an anti-aliasing low-pass filter (45% of sampling frequency), synchronously digitized, and stored locally. We filtered real-time EEG between 0.1-36.0 Hz (2^nd^-order biquad filter), and EMG between 5.0-525.0 Hz (2^nd^-order biquad filter and 40 dB notch filter centred at 50 Hz). These signals informed the detection algorithms for stimuli delivery.

mCLAS was performed as described before^40,49^. In brief, all mice received auditory stimuli consisting of clicks of pink 1/f noise (15ms, 35 dB SPL, 2ms rising/falling slopes) delivered by inbuilt speakers on top of the stimulation chambers. For up-phase-targeted mCLAS, a nonlinear classifier defined four decision nodes comprising power in EEG (EEGpower), power in EMG (EMGpower), a frequency component and a phase-target, which needed to be simultaneously met to deliver sound stimuli, with a trigger delay of ∼[29, 30]ms. The detailed computation of each node is described elsewhere^40,68^. We tracked a 2Hz frequency component within the ongoing EEG signal of both mouse strains, with stimuli targeting either the 30⁰ or 40⁰ phase of ongoing slow waves in the AD (WT and AD mice) or the PD strain (WT and PD mice), respectively^40,49^. We controlled for stimuli-evoked arousals or reflexes via in-built infrared recording cameras in each recording chamber. All triggers were flagged and saved for offline analysis. We recorded 2 EEG/EMG baseline days (no sound stimuli delivered but triggers flagged) and 2 EEG/EMG stimulation days for all mice.

Regarding offline sleep scoring, we scored all recorded files using SPINDLE for animal sleep data^77^, previously validated in our lab for mice^10,40^. SPINDLE retrieved 3 vigilance states - wakefulness (WAKE), non-rapid eye movement sleep (NREM sleep), and rapid eye movement sleep (REM sleep) – from EEG/EMG recordings saved into EDF files, with 4s epoch resolution. WAKE was defined by high or phasic EMG activity and low amplitude but high frequency EEG for more than 50% of the epoch duration. NREM sleep was characterized by the presence of slow waves, reduced EMG activity and increased EEG power in the delta frequency band (0.5-4Hz). REM sleep was hallmarked by high theta power (4-10 Hz) and low EMG activity.

Finally, we pre-processed EEG data offline with a custom-written MATLAB script, as previously described^40,49^. Briefly, we removed artifacts by detecting clipping events, followed by a 3-point moving average and a basic Fermi window function, preventing ringing artifacts^78^. Finally, we resampled the signal at 300Hz and filtered between 0.5–4Hz using low and high-pass zero-phased equiripple FIR filters^68,73^.

### EEG/EMG offline post-processing

#### Slow-wave (SW) detection and characterisation

To infer on SW characteristics upon acute mCLAS in both neurodegeneration models, we implemented a custom-written script in MATLAB (ver. R2023b) adapting several slow-wave detection and characterization pipelines previously described^20,42,52,53,73^. In brief, scored NREM sleep was retrieved for both light and dark periods separately (across 24h recordings), with all positive-to-negative zero crossings, local minimum and maximum between each two consecutive crossings marked and saved for further analysis^73^. These events were considered SWs if: 1) the interval between two consecutive positive-to-negative zero crossings was between 0.2 – 2 s^42,73^; and 2) the minimum amplitude in each of these intervals was greater than 5 µV, to prevent inclusion of random signal fluctuations of low amplitude around 0 µV^20,53^. We did not implement other adaptive amplitude criteria for SW detection^73^, as stricter amplitude thresholds prevent the *a priori* inclusion of shallower SWs present during NREM sleep^20^.

To explore the potential effect of mCLAS on different SW synchronization processes previously described in humans^19,20^, we characterized SWs into different types based on two cut-off thresholds for the amplitude of each SW negative peak:

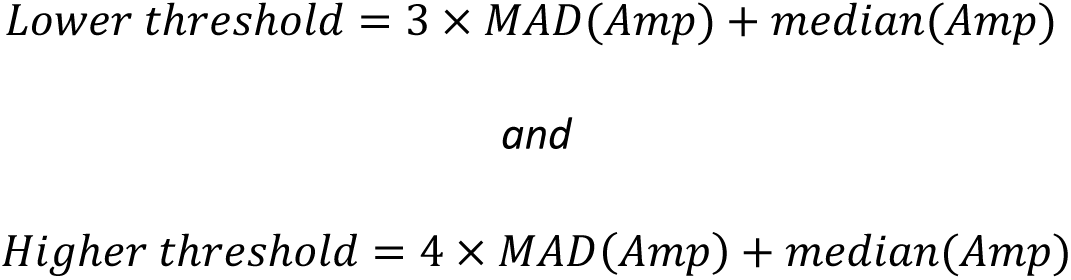

with *MAD* being the median absolute deviation and *Amp* the amplitude of the negative peak.

If the amplitude of a negative peak was below the lower threshold, the SW was considered to be type II, whereas if the amplitude was above the higher threshold, it was characterized as type I; SWs with negative peak amplitudes falling between the thresholds were classified as type III. SW types were classified using negative peak amplitude as the primary metric, as scalp involvement could not be assessed in our single-electrode setup (see Discussion for limitations).

Finally, for each subject and recording day, we calculated both a *Lower* and a *Higher threshold* during each light/dark period. To ensure that stimulation effects were not masked by altered signal properties, these thresholds were determined using baseline data from corresponding light/dark cycles^70^. Specifically, baseline thresholds were first computed for each light/dark cycle using data from two baseline days; original thresholds were then calculated for each stimulation day and stored separately. To select the appropriate threshold for each stimulation day, the two baseline thresholds were compared: if 1) the two baseline thresholds differed by more than 1.5 times (i.e., one was 1.5x larger/smaller than the other), the baseline threshold closest to the original threshold of the stimulation day was chosen; 2) the two baseline thresholds were similar (i.e., differed by less than 1.5 times), their mean was calculated and used as the threshold for that stimulation day’s light/dark cycle.

To characterise each SW type (I, II and III) at baseline and stimulation days across experimental groups and light/dark periods, we assessed the following SW parameters: absolute number of SWs; average duration of a SW, i.e. SW wavelength, defined as the time between two consecutive positive-to-negative zero crossings, with each SW time locked to its negative half wave peak; average amplitude of positive (maximum) peaks, and steepness of ascending slopes from wave trough to wave crest. Finally, we analysed the percentage of change from baseline values of each measure and plotted its 24h dynamics.

#### Sleep spindle detection and characterisation

To compute sleep spindle characteristics, we implemented a custom-written script in MATLAB (ver. R2023b) adapting previously described detection methods^42,55,73^. After the pre-processing and resampling steps, we band-passed filtered the EEG signal with FIR filters between 10-15 Hz (all spindle band), 10-12 Hz (slow spindle band) and 12-15 Hz (fast spindle band). Subsequently, we rectified each signal (root mean squared with a time resolution of 0.003s within a 0.1s time window), and interpolated all maxima via cubic spline interpolation, to create an envelope of each frequency band^55,73^. Only NREM sleep epochs were selected, with each signal divided into light/dark periods for further analysis. We identified sleep spindles within each frequency band as previously described^42,55,73^: when each envelope exceeded an individual threshold within a [0.4, 3]s time window, the event was considered a spindle. We defined individual thresholds by the standard deviation of the filtered signals during all NREM sleep epochs of an individual mouse per day and per light/dark periods, multiplied by a factor of 1.5 (>75 percentile), with the positive and negative threshold crossings corresponding to the onset and offset of a spindle event, respectively. Thresholds of baseline and stimulation days were calculated in this fashion and implemented as described above for SW detection. We characterised all, slow and fast spindles by their onset, calculating spindle density, average spindle duration and average amplitude at spindle peak, for baseline and stimulation days across experimental groups and light/dark periods. Finally, we analysed the percentage of change from baseline values of each measure and plotted its 24h dynamics.

#### SW-spindle coupling event detection and characterisation

After detection of type I, II and III SWs and all, slow and fast spindles, we considered events of SW-spindle coupling when spindle onset co-occurred with the duration of a SW, starting between its onset and offset^45^. Apart from quantifying the number of spindles coupling at the SW peak for baseline and stimulation days across experimental groups, light/dark periods and SW/spindle combinations, we computed the modulation index (MI) and phase slope index (PSI) as measures of cross-frequency coupling strength and directionality, respectively^9,18,54,79,80^. MI is a measure of circular spread indicating how far an empirical distribution deviates from uniformity using the Kullback–Leibler divergence (a high MI signals closely coupling phases around the preferred phase, suggesting a strong coupling) whereas PSI measures the consistency of phase lag or lead between two frequencies (a positive PSI indicates that SWs drive spindles, while a negative PSI indicates that spindles drive SWs)^79,80^. In brief, we extracted 3s data snippets from unfiltered EEG signal corresponding to each SW-spindle coupling event detected and centred at the SW trough. For MI/PSI, cross-frequency coupling strength/directionality were calculated between the phase of a lower frequency (SWs, 0.5–4 Hz in 0.29/0.5 Hz steps for MI/PSI, respectively) and the amplitude of a higher frequency (sleep spindles, 10–15 Hz in 1 Hz steps). We defined a sliding window of 1s with 0.5s steps, using six SW cycles to reliably estimate frequency power^80^. Subsequently, we averaged MI and PSI values obtained for all frequency sub-band pairs to obtain a single estimate for MI and PSI at baseline and stimulation days across experimental groups, light/dark periods and SW/spindle combinations.

Additionally, we performed event-locked spectral analysis of SW-spindle coupling events adapting previous pipelines implemented on EEG sleep data of older adults and MCI patients^9,55^. Briefly, we calculated time-frequency representations (TFRs) for each 3s SW-spindle coupling snippet centred in the SW trough and filtered in the delta band (0.5-4 Hz, low and high-pass zero-phased equiripple FIR filters). For each snippet, trials with excessive broadband amplitude were excluded using a robust RMS-based threshold (>6× median RMS). To minimize edge artifacts in the time-frequency transform, we applied ∼100 ms of reflection padding to each trial before detrending. We obtained spectral estimates between 5–20 Hz (in 0.25 Hz steps) using a sliding Hanning tapered window (10ms steps) with a variable frequency-dependent length^55^, including six cycles to ensure reliability in power estimates as described previously. For each experimental group and SW/spindle combination, power estimates were aggregated using the median across trials to improve robustness against outliers, and baseline correction was achieved by subtracting the mean power in a −1.5 to −0.5s pre-event window. To preserve temporal structure while reducing spectral noise, we applied a Savitzky–Golay filter (5-point, second-order) along the frequency domain only, with TFRs subsequently cropped to a peri-event interval of −0.5 to +0.5 s. In parallel, we extracted the grand average EEG signal (mean ± SEM) of all SW events locked to the SW trough (t=0s) for each experimental group and SW/spindle combination^55^. To illustrate stimulation- and genotype-related changes, we additionally computed difference spectrograms between predefined contrasts (e.g WT vs AD, baseline vs stimulation)^55^. We computed difference SW traces by subtracting the group-mean waveforms, with the variance of the SEMs combined in quadrature. Finally, we extracted mean spindle (10–15 Hz) spectral power at its maximum, SW trough and peak time points, detected from the group-level average SW trace corresponding to each SW-spindle combination and experimental group, for baseline and stimulation days across light/dark periods.

#### Slow-wave energy (SWE) characterisation

As a correlate of sleep pressure dissipation and its homeostatic regulation, we computed slow-wave energy (SWE) as the cumulative sum of delta power in NREM sleep in each 24h recording, for baseline and stimulation days across experimental groups^10,23^.

### Statistics

Raw data were processed in MATLAB 2023b (Mathworks, USA). We conducted statistical analysis in IBM SPSS Statistics 28 (IBM, USA), with data plotted in GraphPad Prism 11.0.0.84 (GraphPad Software, LLC) unless stated otherwise. We inspected the data for outliers through boxplots, with outliers considered as values 1.5 box-lengths from the edge of the box.

We assessed the distribution of each variable and its residuals for normality through skewness and kurtosis, while the homogeneity of variances was verified using Levene’s test with 5% significance. If normality and homogeneity of variances were met, one sample t-test, two-sample t-test, one-way ANOVA with Holm-Šidák post hoc test, or a two-way mixed ANOVA (experimental group*time) with Holm-Šidák post hoc test were applied. If the Mauchly’s test of sphericity indicated that the assumption of sphericity was violated for the two-way interaction, we interpreted the results with the Greenhouse-Geisser correction. Each test is reported with the corresponding p-value, unadjusted in case of single comparisons or adjusted in case of multiple comparisons, with significance set to 0.05. Results are shown as the mean and SEM.

## Data and code availability

All data generated or analysed during this study are included in the manuscript and supporting files and/or available upon request.

## Acknowledgments and funding statement

This project was funded by Parkinson Schweiz (ID, DN), the Dementia Research Switzerland-Synapsis Foundation (2018-PI03) via an earmarked donation of the Armin and Jeannine Kurz Stiftung (DN), the Hurka Foundation (DN) and the Swiss National Science Foundation (MAZ, Grant Nr. 215333; CRB, Grant Nr. 188790; DN, Grant Nr. 215046). The funding sources had no involvement in study design, collection, analysis, interpretation of the data nor in writing or deciding to submit this article for publication.

## Author contributions

I.D.: Conceptualization; methodology; software; validation; formal analysis; investigation; data curation; visualization; funding acquisition; writing – original draft; writing – review and editing. M.A.Z.: Software; writing – review and editing. C.R.B.: Funding acquisition; writing – review and editing. D.N.: Conceptualization; resources; funding acquisition; supervision; project administration; writing – original draft; writing – review and editing.

