## Supplementary material for "Bidirectional restoration of sleep homeostasis in neurodegeneration via closed-loop auditory stimulation": Suppl. Figures 1-3

**Containing:**

Figures S1 to S3

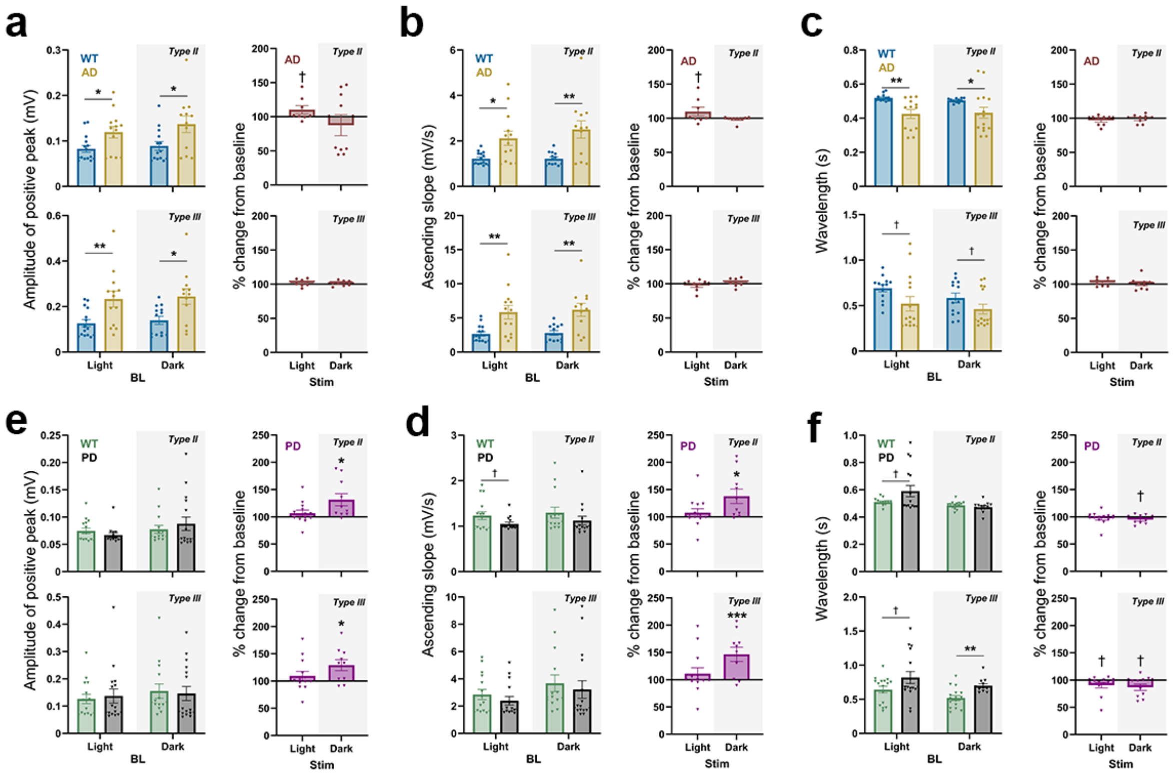


**Figure S1 – Type II and III SW characteristics in AD and PD mice. a, b and c)** Left panels show the positive peak amplitude, ascending slope or wavelength of type II or III SWs at baseline (BL) for WT and AD mice. Right panels represent the same measures as percentage of change from BL during light and dark periods in AD mice (Stim); **d, e and f)** Left panels show the positive peak amplitude, ascending slope or wavelength of type II or III SWs at BL, for WT and PD mice. Right panels represent the same measures as percentage of change from baseline during light and dark periods in PD mice (Stim). In the bar plots, each scatter dot represents average daily values of each animal, each bar the mean, and the whiskers the SEM. The grey shaded panel corresponds to the dark period. ***p<0.001, **p<0.01, *p<0.05, †<0.2.

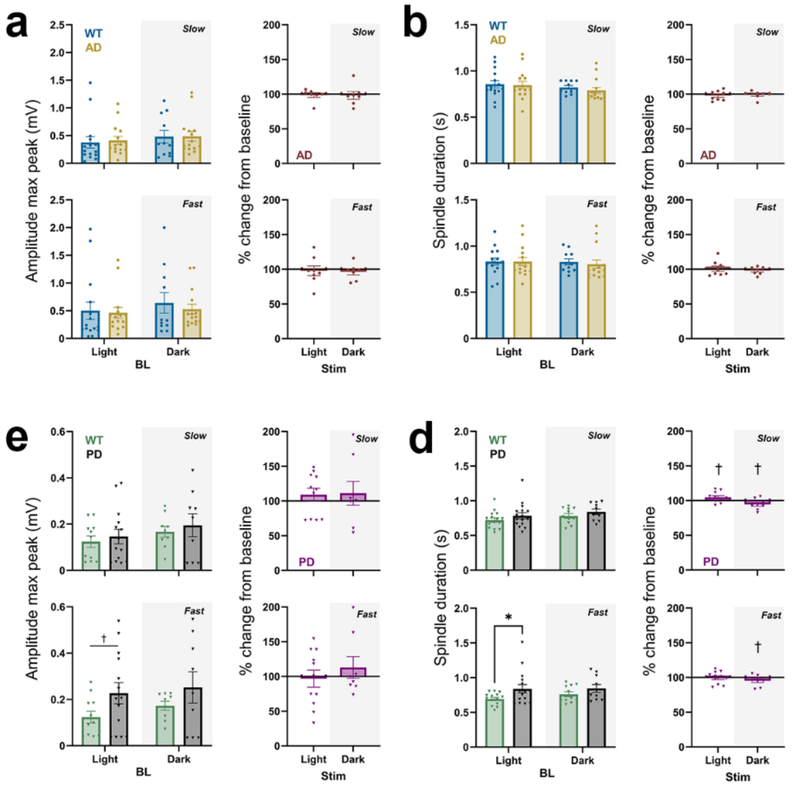


**Figure S2 - Sleep spindle modulation by mCLAS in AD and PD mice. a and b)** Left panels show the amplitude of maximum peak and spindle duration of slow or fast spindles at baseline (BL) for WT and AD mice. Right panels represent the same measures as percentage of change from BL during light and dark periods in AD mice (Stim); **c and d)** Left panels show the amplitude of maximum peak and spindle duration of slow or fast spindles at baseline (BL) for WT and PD mice. Right panels represent the same measures as percentage of change from BL during light and dark periods in PD mice (Stim). In the bar plots, each scatter dot represents average daily values of each animal, each bar the mean, and the whiskers the SEM. The grey shaded panel corresponds to the dark period. *p<0.05, †<0.2.

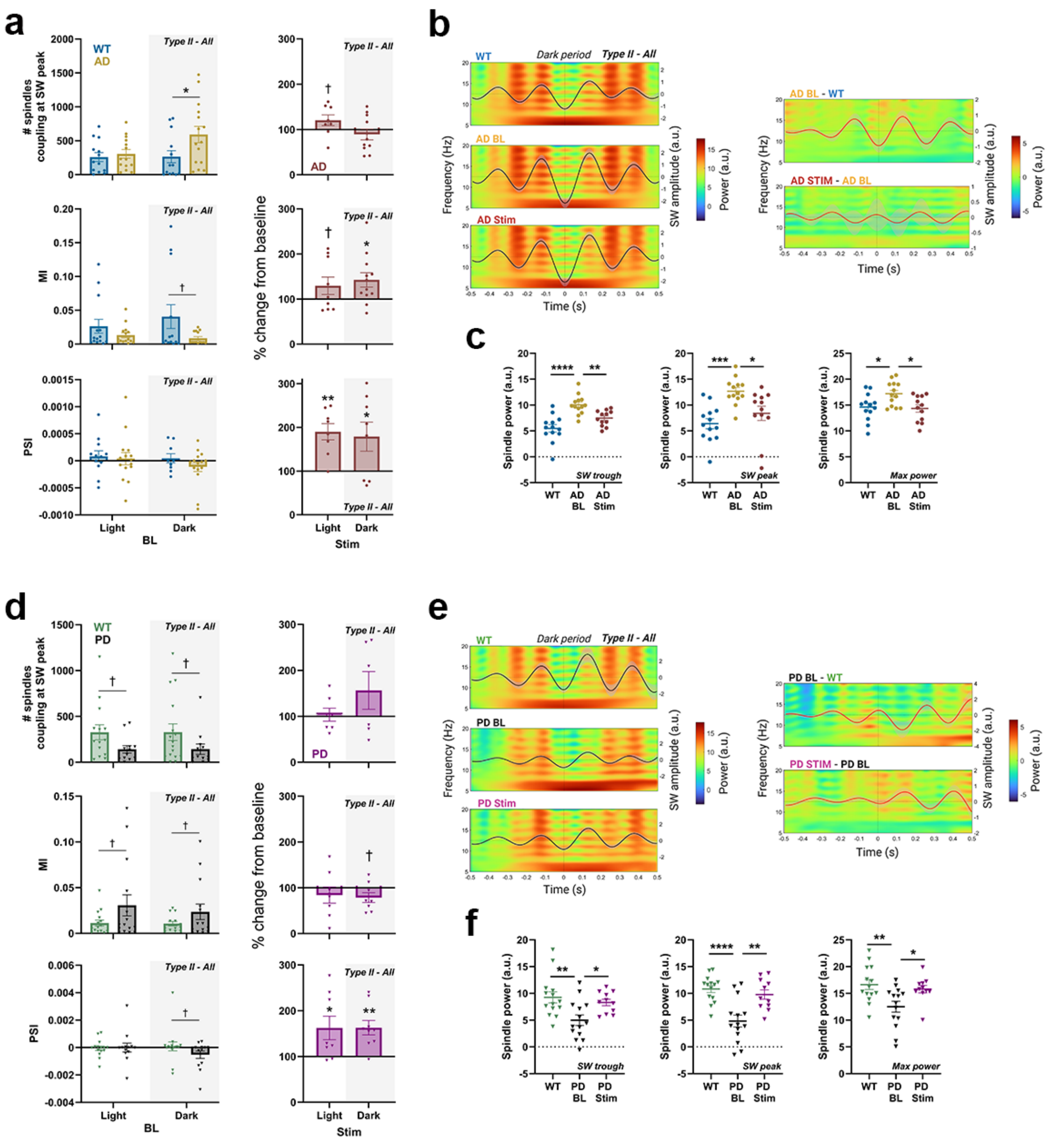

**Figure S3 – Type II SW - all spindle modulation by mCLAS in AD and PD mice. a)** Left panels show either the number of spindles coupling at SW peak (top), MI (middle) or PSI (bottom), for SWs type II – all spindles coupling at baseline (BL) in WT and AD mice in the light and dark periods; Right panels show the percentage of change from BL during light and dark periods in AD mice (Stim) across those measures; **b)** Left panels show TFRs and average SWs traces for type II – all spindles coupling during the dark period in WT, AD mice at baseline (AD BL) and during stimulation days (AD stim). Right panels show difference TFRs and average SW traces between AD BL-WT and AD Stim-AD BL; **c)** Spindle power at SW trough, SW peak and maximum spindle power extracted from TFRs in b for WT, AD BL and AD Stim; **d)** as in **a** but for PD mice; **e)** as in **b** but for PD mice; **f)** as in **c** but for PD mice. In the bar and scatter plots, each scatter dot represents average daily values of each animal, each bar the mean, the whiskers the SEM and the grey shaded panel corresponds to the dark period. In the TFR plots, black solid lines correspond to the average SW, red solid lines correspond to the average SW difference, shaded areas correspond to the SEM and the dotted line at t = 0 seconds correspond to the SW trough (event center). ***p<0.001, **p<0.01, *p<0.05, †<0.2. Schematics in a and b were created with BioRender (https://BioRender.com). TFRs in c and f were created in MATLAB (R2023b).
